# Antimicrobial peptides play a functional role in bumblebee anti-trypanosome defense

**DOI:** 10.1101/010413

**Authors:** Soni Deshwal, Eamonn B. Mallon

**Affiliations:** Department of Biology, University of Leicester

**Keywords:** *Crithidia bombi*, antimicrobial peptides, RNAi, gut microbiota

## Abstract

Bumblebees, amongst the most important of pollinators, are under enormous population pressures. One of these is disease. The bumblebee and its gut trypanosome *Crithidia bombi* are one of the fundamental models of ecological immunology. Although there is previous evidence of increased immune gene expression upon *Crithidia* infection, recent work has focussed on the bumblebee’s gut microbiota. Here, by knocking down gene expression using RNAi, we show for the first time that antimicrobial peptides (AMPs) have a functional role in anti-*Crithidia* defense.

## INTRODUCTION

Bumblebees are among the most important wild and agricultural pollinators with, for example, 25 major crops grown within the E.U. pollinated by bumblebees (Cameron et al., 2011; Corbet et al., 1991). Throughout the northern hemisphere, bumblebee populations have being decreasing precipitously (Cameron et al., 2011; Goulson et al., 2005). Although this is probably the result of multiple factors, a key component is disease (Cameron et al., 2011).

The bumblebee (*Bombus terrestris*) is an annual, eusocial insect and host to a variety of parasites including the trypanosomatid, *Crithidia bombi* (Trypanosomatidae, Zoomastigophorea) (Lipa and Triggiani, 1980). The bumblebee/*Crithidia* system is one of the main models for the study of ecological immunology (Schmid-Hempel, 2001). *C. bombi* colonizes the bumblebee’s gut and begins releasing transmission stages into the faeces 2–3 days after infection. Parasitemia peaks around 7–10 days into the infection (Schmid-Hempel and Schmid-Hempel, 1993). *C. bombi* prevalence in *B. terrestris* populations ranges from 10–30% but can be much higher (80%) (Shykoff and Schmid-Hempel, 1991) with increased inter-colony transmission as the season progresses (Imhoof and Schmid-Hempel, 1999). A variety of fitness effects are seen at the colony level due to infection: queens have reduced success in colony founding (Brown et al., 2003), colonies have smaller worker populations and produce fewer sexual offspring (Brown et al., 2000), and the ability to learn floral cues is impaired in infected workers (Gegear et al., 2006). The virulence (i.e. parasite induced host-death) of the parasite is condition-dependent. For example, mortality rates of infected bumblebees increases by up to 50% after 10–15 h under starvation conditions, relative to non-infected, starved controls (Brown et al., 2000).

Compared to the extensive ecological understanding we have of the *B. terrestris* - *C. bombi system*, its dynamics at the molecular level are only beginning to be revealed. Standing levels of prophenoloxidase increase upon infection with *C. bombi* (Brown et al., 2003a). Several quantitative trait loci are linked to immune defence against this parasite (Wilfert et al., 2007). Four studies have looked at differences in bumblebee gene expression upon infection (Brunner et al., 2013; Riddell et al., 2009, 2011; Schlüns et al., 2010). Chief among the genes found have been antimicrobial peptides (AMPs).

As well as the above evidence of AMP expression in response to *Crithidia* infections, there is growing evidence of expression of AMPs and their efficacy in response to medically important trypanosomes in vector insects (McGwire and Kulkarni, 2010). AMPs attack trypanosomes by numerous mechanism with most focussed on the cell membranes of trypanosomes (Harrington, 2011; McGwire and Kulkarni, 2010; McGwire et al., 2003).

However the role of the immune system in *Crithidia* defense has been called into question. Recently it has been shown that gut microbiota has a central role in the bumblebee’s response to *Crithidia* (Koch and Schmid-Hempel, 2011). Although there is data on transcription supporting a role for AMPs in *Crithidia* defense, they have never been shown to have a functional role. Along with the confusion of the role of the immune system in anti-*Crithidia* defense generally (Hauton and Smith, 2007; Koch and Schmid-Hempel, 2011), this presents a need for a clear display of the functional role of AMPs in response to *Crithidia*. Here, we knocked down *Defensin* and *Abaecin* expression and show that bumblebees thus treated, are less able to defend themselves against *Crithidia* compared to controls.

## MATERIAL AND METHODS

### Bumblebees and *Crithidia*

A *B. terrestris* colony was obtained from Kopert Biosystems UK. Bumblebees were maintained in standard conditions: 27°C, 60% humidity under red light. They were fed 50% meliose solution and pollen *ad libitum*. *Crithidia bombi* was collected from wild queens of *B. terrestris*. Queens were captured from the botanical gardens of the University of Leicester in March 2013. Faeces was collected and examined for presence of *C. bombi* cells. Ten colony workers were infected with a mixture of faeces and meliose to produce C. *bombi* cells for further experiment.

### RNA interference

The sequences of *Defensin* (FJ161699.1), *Abaecin* (GU233780) and the *Nautilus* (X56161.1) genes were retrieved from NCBI. Drosophila *Nautilus* shows no similarity to the genes of B. terrestris and hence is used as a RNAi control. All the primers were designed using Primer3 (version 0.4.0). To amplify *Defensin* and *Abaecin*, DNA was extracted from bumblebee using Qiagen DNA mini kit. For *Nautilus*, w118 flies were kindly provided by Dr. Ezio Rosato, Department of Genetics, University of Leicester. Polymerase chain reaction (PCR) was performed using 1 microlitre of DNA template, 200 nM of each primer, 2.5 mM of MgCl2, and York Bioscience Taq DNA Polymerase (dNTPs and buffer included) in a 25 microlitre reaction using PCR conditions 95°C for 3 min and 35 cycles of 95°C for 30 s denaturation, 30 s annealing, 72°C for 30 s extension. The annealing temperature was varied according to the primers used (Table S1). Amplified products were checked for size on 1% agarose gel and bands extracted using Wizard^®^ SV Gel and PCR Clean-Up System.

T7 promoter sequence 5’-TAATACGACTCACTATAGGGAGA-3’ was added to the purified amplification products of the genes (Defensin and Nautilus) using PCR to generate a transcription template. The primers were synthesized by adding T7 promoter sequence to the 5’ end of the gene specific primers. One hundred nM T7 promoter-containing PCR primers (sense and antisense) were used in a single PCR reaction to generate transcription template for both strands of the dsRNA. Since, the first cycles of PCR used only the 3’ half of the primers, the gene-specific part, the annealing temperature for the first 5 PCR cycles was 5°C higher than the calculated Tm for the gene-specific region of the primer. After 5 cycles, the annealing temperature for the entire primer (Table S1) was used for the following cycles: 95°C for 30 s denaturation, 30 s annealing and 72°C for 30 s extension for 35 cycles. The bands were visualized and extracted as above. 1 microgram of DNA with T7 promoter overhangs was used to construct dsRNA, in vitro, using MEGAscript^®^ T7 Kit (Invitrogen) as per manufacturer’s instructions. The concentration of dsRNA (RNAi) was determined at 260 nm using a Nanodrop ND-1000 Spectrophotometer.

### RNAi injection and *C. bombi* infection

Thirty bees in total were used in this experiment. Twenty seven bees were injected (16.6 microlitre of elution buffer containing 1 microgram of dsRNA) with either RNAi *Defensin* (11) RNAi *Abaecin* (5) or RNAi *Nautilus* (11). Five bees were used for cell counts (each RNAi group) and six for qPCR (RNAi Defensin and RNAi Nautilus only). Three uninfected and elution buffer injected bees was used as control samples for qPCR.

After injections, bees were kept separately and fed with 50% diluted nectar and pollen for 24 h. Bees were then fed 20 microlitre of a 1000 cells/microlitre *Crithidia* inoculum (8 bees each for infected RNAi *Defensin* and RNAi *Nautilus* groups and 5 bees for RNAi *Abaecin*) or 20 microlitre of meliose solution (3 bees each for uninfected RNAi Defensin and RNAi Nautilus groups only). Bees used for cell counts were left for a further 5 days. *Crithidia* cells were counted in 10 microlitre samples of each bee’s faeces using a haemocytometer. Bees used for qPCR were sacrificed 24 hours after infection (Riddell et al., 2011).

### Quantatitive PCR (qPCR)

Total RNA was extracted from whole bees using RNAeasy mini kit (Qiagen). cDNA was synthesized from RNA template by using Tetro cDNA Synthesis kit (Bioline). One microgram of RNA template, 10 mM of dNTP mix, 5X RT buffer, Ribosafe RNAse inhibitor 2 microlitre and tetro Reverse Transciptase 2 microlitre were used and the reaction volume was made up to 40 microlitre with DEPC water. A combination of OligodT and Random Hexamers primers (1 microlitre each) was used to improve the reverse transcription efficiency of RNA templates. The prepared samples were incubated for 10 min at 25°C followed by 45°C for 30 min. The reaction was terminated by incubating samples at 85°C for 5 min.

Two microlitre of cDNA was added to a qPCR reaction mix containing 10 microlitre 2X SYBR Green JumpStart Taq ReadyMix (Sigma-Aldrich, UK) 7.6 microlitre ddH2O and 0.4 microlitre of the forward and reverse primer mix (5 micromoles each). Two sets of primers were used, Defensin, and Actin (XM 003394442) (Table S1). Actin was used as an endogenous control (reference gene). Actin has previously been used in this role in gene expression studies of bumblebee immunology (You et al., 2010). For each sample three technical replicates were run. qPCR was carried out on a PTC-200 MJ research machine thermal cycler with a Chromo 4 continuous fluorescence detector, using standard settings and Opticon software v4.7.97.A for Windows 2000 Professional. The qPCR was carried out using protocol: 95°C for 4 min followed by 40 cycles of 30 s for 95°C denaturation, 30 s for 60°C annealing and 30 s products were checked products were checked products were checked products were checked products were checked for 72°C extension. The relative expression ratio (Pfaffl, 2001) was calculated as 

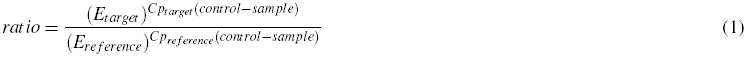

where E is qPCR efficiency and Cp is the crossing point of each transcript, target refers to the target gene, reference is *Actin*, and control is the uninfected, elution buffer injected samples.

## RESULTS AND DISCUSSION

There was a significant difference between treatments in levels of *Crithidia* infections (F_2,12_ = 28.95, p *<*0.0001). RNAi *Defensin* treated bees had higher levels of *Crithidia* infection compared to RNAi Nautilus controls (Tukey HSD post-hoc test: p = 0.0008). RNAi Abaecin bumblebees also have higher levels of Crithidia infection compared to RNAi Nautilus controls (Tukey HSD post-hoc test: p *<*0.0001). See Figure 1.

**Figure 1.**
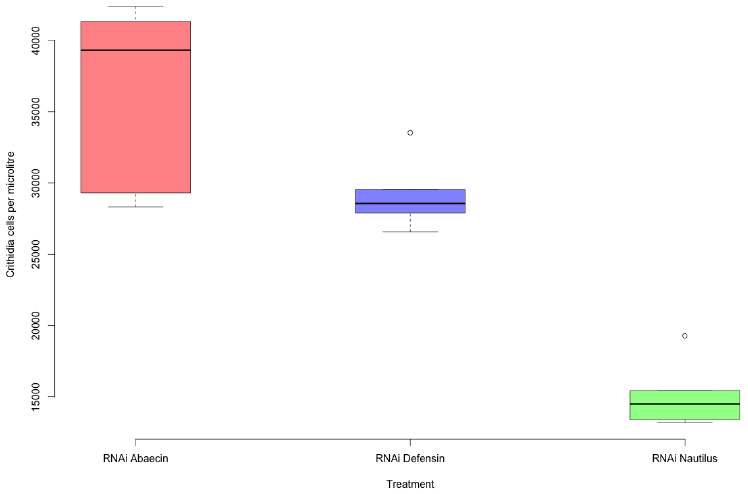
Boxplots of levels of Crithidia infection for various RNAi treated bees.

To confirm that it was actually decreased immune gene expression that caused this increased parasitemia, we measured *Defensin* expression in our RNAi *Defensin* vs. RNAi *Nautilus* samples. Treatment had a significant effect on Defensin expression levels (F_1,8_ = 14.27, p = 0.0054). RNAi *Defensin* treated bees had lower levels of Defensin expression relative to RNAi *Nautilus* treated controls. This is true in both infected and uninfected groups. Infected RNAi *Defensin* treated bees had similar levels of *Defensin* expression as uninfected controls (RNAi *Nautilus*). See Figure 2. As would be expected (**?**), infection caused a significant increase in *Defensin* expression (F_1,8_ = 11.896, p = 0.00871) for both RNAi *Defensin* and RNAi *Nautilus* groups.

**Figure 2.**
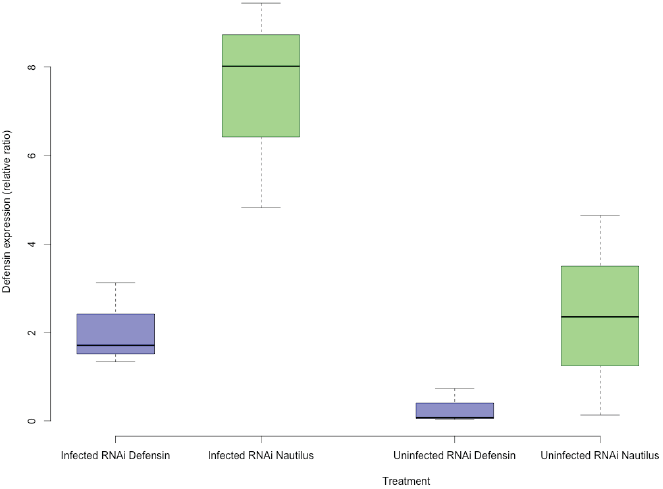
Boxplots of *Defensin* expression for infected and uninfected RNAi *Defensin* and RNAi *Nautilus* (control) treated bees.

We have shown that by knocking down AMP expression we increase the infection levels of *Crithidia*. This is evidence for the central role of the bumblebee immune system in the fight against *Crithidia*. This seems to contradict recent findings that found a central role for gut microbiota in the bumblebee’s defense against *Crithidia* (Koch and Schmid-Hempel, 2011). It is clear that the outcome of a *Crithidia* infection is not the result of the immune system or the gut microbiota but rather the interaction between the parasite, the gut microbiota and the host immune system (Castro et al., 2012; Garcia et al., 2009; Weiss et al., 2013). This interaction will be the focus of much future research.

## CONCLUSIONS

We found that reducing expression of two AMPs (Defensin and Abaecin) lead to an increase parasitemia of *Crithidia bombi* in bumblebees. This is strong evidence that AMPs are powerful anti-trypanosome agents.

